# Birds protected in Europe, overlooked in Africa: conservation synergies between African primates and wintering Afro-Palearctic migratory birds

**DOI:** 10.64898/2026.07.20.739569

**Authors:** Issaka Abdou Razakou Kiribou, Jean Pierre Samedi Mucyo, Paul K. Kazaba, Solange Mekuate Kamga, Hjalmar S. Kühl, Jessica Junker, Tim Sasse, Stefanie Heinicke

## Abstract

Primates are frequent targets of international conservation efforts, due to their anthropological significance and function as biodiversity conservation flagship species. Afro-Palearctic migratory birds receive considerable conservation attention in their breeding habitats, but much less in their wintering areas. Good wintering habitat conditions are, however, crucial for long-term population viability. Integrating complementary conservation efforts for these taxa may provide substantial co-benefits. Here, we evaluate the spatial, ecological, and conservation congruence between 248 Afro-Palearctic migratory bird and 213 African primate species to identify potential synergies in intercontinental conservation efforts. The wintering areas of 218 migratory bird species overlap with primate ranges. Up to 125 Afro-Palearctic migratory bird species co-occur with at least one primate species locally and highest regional cross-taxon richness occurs in forest-savanna ecotones in West Africa and eastern DRC. Agriculture and hunting threaten both taxa (88-97% of primate and 59-70% of bird species). Several conservation interventions have demonstrated effectiveness for both taxa, particularly habitat protection and hunting reduction. We propose three policy priorities: strengthening cross-taxa monitoring within existing primate research infrastructure, improving the conservation evidence base in the African context, and creating financing synergies that allow conservation investments to simultaneously benefit both taxa.

## 1 Introduction

As threats to biodiversity continue to increase, migratory birds are particularly vulnerable and often have stronger population declines than resident species (Bairlein 2016). This is because, throughout their life cycle, migratory species encounter diverse threats across different regions, including habitat loss, hunting pressure, and the impacts of climate change (Buchan *et al* 2022). Afro-Palearctic migratory birds, i.e., birds that breed in Europe and overwinter in Africa, are a prominent example (Marcacci *et al* 2023). Although birds are a major focus of conservation activities across Europe, populations continue to decline, and trends are more pronounced in long-distance migrants compared to resident species (Burns *et al* 2021). One reason is that they receive disproportionately little conservation attention across their overwintering areas across sub-Saharan Africa (Marcacci *et al* 2023), even though environmental conditions during the non-breeding period strongly influence survival during spring migration and breeding (Cooper *et al* 2024). In contrast, African primates are important conservation flagship species and a focus of research and conservation activities (Heinicke *et al* 2021). A significant portion of this conservation funding is of European origin. For example, a third of funding for biodiversity conservation in Madagascar comes from European donors (Eklund *et al* 2025). But little of this money is targeted to wintering migratory birds (Marcacci *et al* 2023). While other species might overlap more strongly with wintering birds, they neither serve as flagship species nor receive the conservation funding that primates do.

Both African primates and Afro-Palearctic migratory birds are broadly distributed across sub-Saharan Africa, raising the question of whether their geographic ranges, habitat use, and threats overlap to a degree that would allow shared conservation effort (Fig. 1). Where such overlap exists, conservation investments directed at primate habitat could simultaneously benefit wintering bird populations. For example, the recent establishment of a new national park in West Africa (i.e. Moyen Bafing in Guinea) to protect a chimpanzee population, now protects an area known to harbour 203 bird species (Muench *et al* 2021). However, whether such co-benefits are incidental or could be systematically achieved through integrated conservation planning has not been assessed across both taxa.

**Figure 1.**
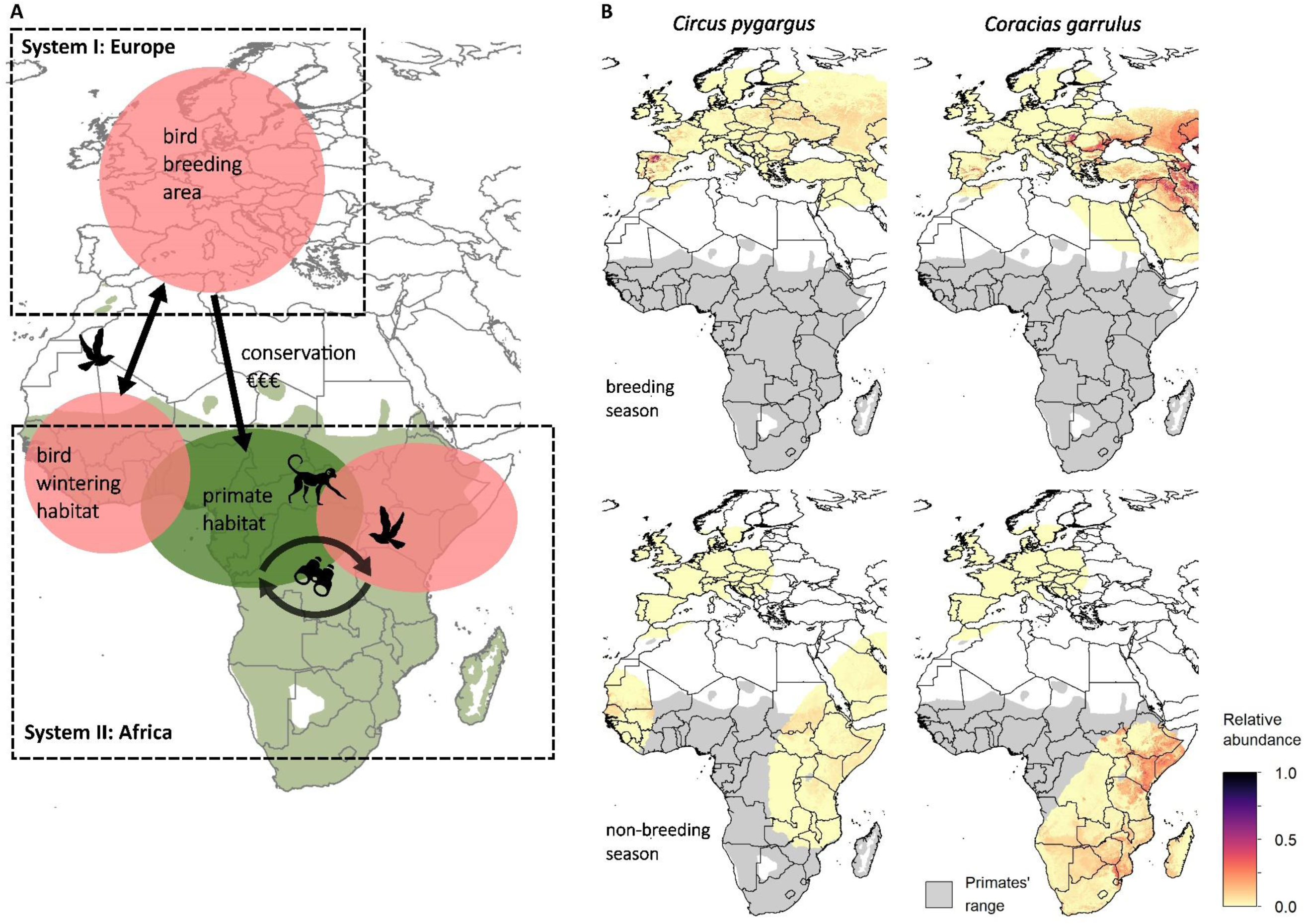
(A) Conceptual framework illustrating ecological and economic linkages between Europe and Africa through migratory bird species that breed across Europe (System I) and overwinter across Africa (System II). Green areas show the geographic range of African primates. (B) Relative abundance maps of breeding (top) and non-breeding (bottom) seasons for two exemplary migratory bird species. Migratory birds receive relatively little conservation attention in overwintering areas, while at the same time African primate species are conservation flagship species that are often targets of international conservation funding. Explicitly considering these intercontinental linkages can help to identify conservation synergies across the two taxa, as well as for sympatric biodiversity (e.g., joint use of field stations). Icons designed by Freepik (www.freepik.com).

Realising conservation synergies between these taxa requires identifying whether the threats they face are similar and whether proven conservation interventions for one taxon are applicable to the other. We use the term conservation synergies to refer to cases where conservation interventions targeting one taxon deliver measurable benefits to the other, and where integrated planning across both groups is therefore more effective than managing each in isolation. The Conservation Evidence Project documents effective conservation interventions for both birds and primates (Williams *et al* 2013, Junker *et al* 2017), enabling an assessment where interventions could effectively serve both taxa. At the same time, European conservation funding flowing into Africa could be strategically extended to also address the non-breeding habitat needs of migratory birds, at relatively low additional cost.

Here, we conducted a range-wide analysis across 213 African primate species and 248 Afro-Palearctic migratory bird species to assess the extent to which their geographic ranges, habitat use, threats, and applicable conservation interventions overlap. From this analysis, we derive specific policy recommendations for conservation practitioners and funders seeking to maximise co-benefits for both taxonomic groups.

## 2 Methods

### 2.1 Species list and species data

To derive a list of bird species that breed in Europe and overwinter in Africa, we began with the 72 species compiled by Beresford *et al* (2019). We added species from the “List of Western Palearctic bird species migrating within Africa” (Walther 2005), which includes birds that are regular breeders within the Western Palearctic and migrate within Africa. From this second list, we excluded all birds that do not occur in Europe and all seabirds not found in terrestrial Europe or Africa (Jonsson 1992). This yielded a final list of 248 migratory bird species.

Geographic range data for all African primate species (n=213) were obtained from the IUCN Red List of Threatened Species (IUCN 2025), and range data for all bird species were obtained from BirdLife International (2023). We created a grid (10 km resolution) using the Lambert azimuthal equal-area projection centered on Africa, and all spatial layers were reprojected accordingly. Given the coarse resolution of species range maps, we complemented the spatial overlap analysis with occurrence records from the Global Biodiversity Information Facility (GBIF) database (Calabuig 2013). We calculated minimum distances between occurrence points of primate-bird species pairs for which the geographic range overlapped by more than 80%. Occurrence data were available for 332 bird-primate pairs. Due to spatial bias and incomplete coverage in the GBIF database, particularly for African species (Beck *et al* 2014), we did not use these data to estimate average inter-species distances. Instead, we used the occurrence-based distance analysis to verify that species pairs identified as overlapping from range maps also show evidence of local co-occurrence, confirming that overlap extends beyond broad-scale range coincidence.

We compiled information on threat status, population trend, habitat, and type of threat from the IUCN Red List of Threatened Species (IUCN 2025). Birds’ relative abundance data used in Figure 1 were retrieved from eBird (Fink *et al* 2024).

### 2.2 Conservation interventions

We include protected areas spatial data designated as ‘national park’ or IUCN categories I and II from the World Database on Protected Areas (UNEP-WCMC and IUCN 2024), and Key Biodiversity Areas (BirdLife International 2025). To compare the conservation interventions listed for birds with those listed for primates and to determine the extent to which they overlap, information on the effectiveness of conservation interventions was compiled from the Conservation Evidence Project (Williams *et al* 2013, Junker *et al* 2017). For the analysis, we retained only those conservation interventions with an effectiveness rated as either *‘beneficial’* or *‘likely to be beneficial’*. Analyses were implemented in R version 4.5 (R Core Team 2025) with the following R packages: ‘classInt’, ‘dplyr’, ‘sf’, ‘shapefiles’, ‘splancs’.

## 3 Results

### 3.1 Spatial overlap and habitat use

Our analysis reveals substantial geographic overlap between wintering Afro-Palearctic migratory birds and African primates (Figure 2, Table S1). The geographic range of 218 of the 248 migratory bird species included overlaps with the geographic range of primates, which adds up to a total area of spatial overlap of more than 19 million km². Only 30 bird species do not overlap with any primate species. Within the geographic range of primates, the number of species per grid cell ranges from 1 to 19 primate species and 0 to 125 migratory bird species. For 119 bird species, at least 30% of their range overlaps with the range of primates. The distance of occurrence points analysis implemented to verify whether species with overlapping geographic ranges also co-occur showed that for 26 bird-primate species combinations, the minimum distance of occurrence points was 0 km. 25% of all bird-primate species combinations analysed here (n=332) have a minimum distance of less than 2 km (Figure S1).

**Figure 2.**
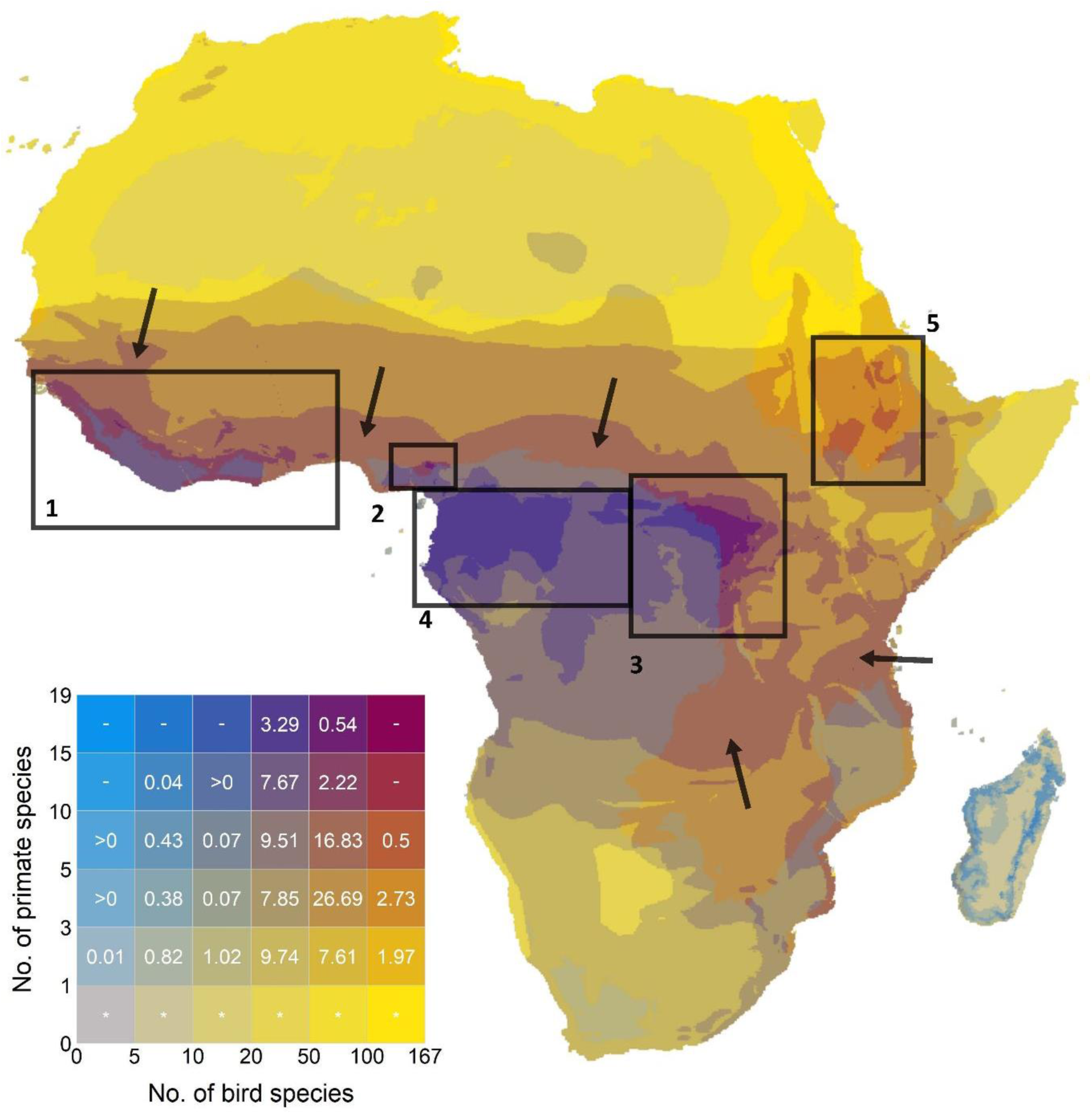
Spatial overlap of African primate and Afro-Palearctic migratory bird species. Coloured boxes in the legend: numbers quantify the proportion of grid cells in each category relative to the number of grid cells within primate range (in %), colours marked with (*) are outside of primate ranges, (-) combination does not occur in the map. Black boxes mark areas of strong overlap, specifically areas of high primate and bird species richness are (1) Guinean forests, (2) Cameroon highland forests, (3) Albertine rift highland forests and Congolian lowland forests; areas of high primate and medium bird species richness are (4) Congolian coastal and lowland forest; areas of medium primate and highest bird species richness are (5) Ethiopian montane grasslands and woodlands, and black arrows mark areas with medium primate and high bird species richness, including Guinean forest-savanna, Sudanian Savanna, Northern Congolian Forest-Savanna and Dry miombo woodlands.

Highest primate species richness can be found in forested areas, for example, forests in the Congo basin (Box 4 in Figure 2, Figure 3B), while bird species richness is highest in more open habitats, such as woodlands and savannas (black arrows in Figure 2, Figure 3C). However, hotspots of overlap in both, bird and primate species richness, are spread across tropical Africa, including the Guinean forests in West Africa, Congolian coastal and lowland forest in Western Equatorial Africa, and the Albertine rift highland forests in East Africa (Figure 3). Notably, areas at the ecotone of forest and savanna areas, as well as woodlands, are areas with medium primate richness (i.e., 5-9 species) and high bird species richness (i.e., at least 50 species), specifically the Guinean and Congolian forest-savanna, Sudanian savanna and Miombo woodlands in southern Africa (Figure 3C, Figure 4).

**Figure 3.**
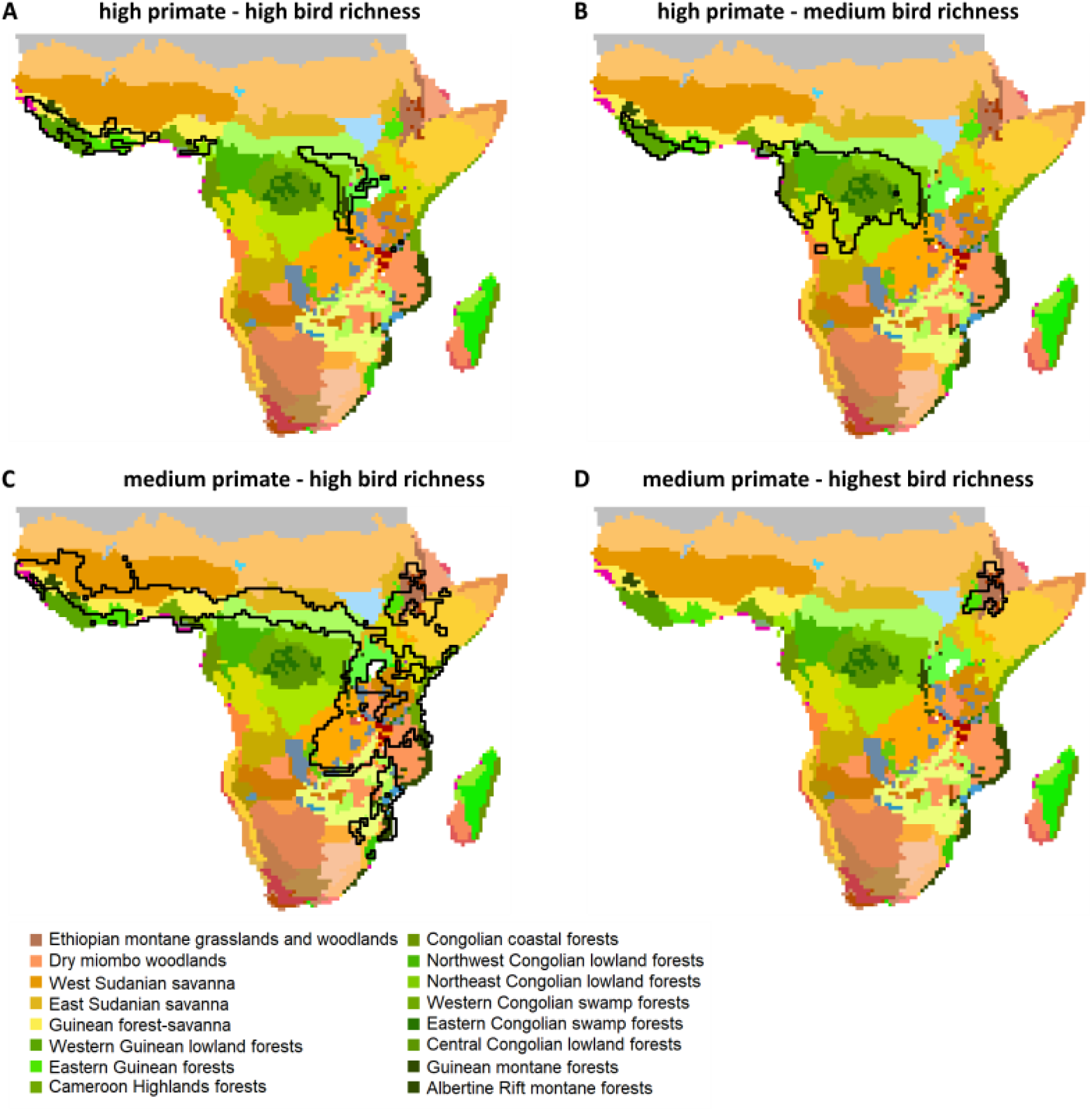
Areas of medium to high bird and primate species richness overlayed to African ecoregions. (Dinerstein *et al* 2017). Black outlines highlight areas with grid cells with A) ≥ 50 bird and ≥ 10 primate species (corresponding to dark red areas in Fig. 4), B) ≥ 20 bird and ≥ 10 primate species, C) ≥ 50 bird and ≥ 5 primate species, D) ≥ 100 bird and ≥ 5 primate species.

**Figure 4.**
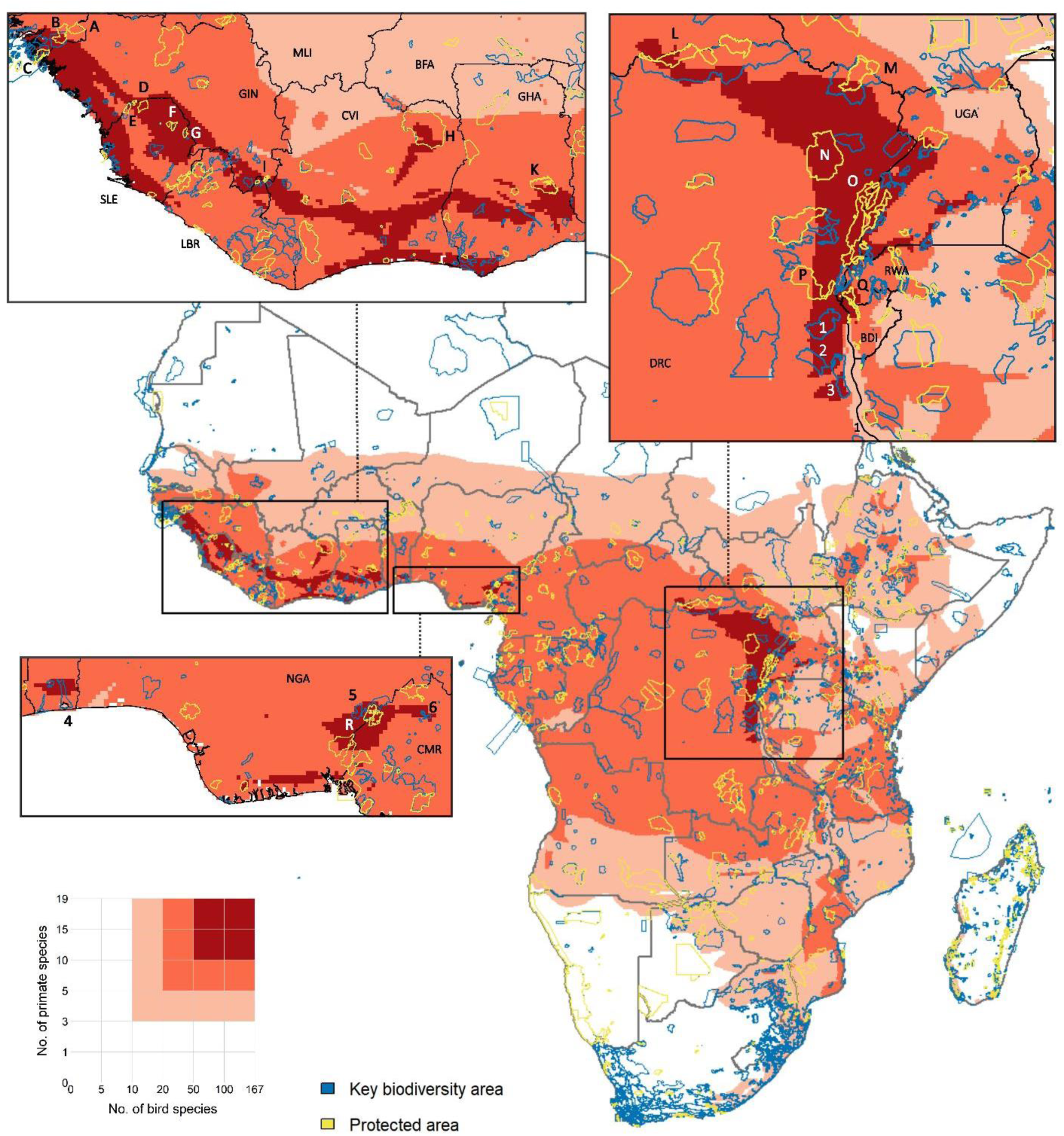
Map highlighting the areas with highest species richness in primates and migratory birds. Light-red: grid cells with ≥ 10 bird and 3 primate species, mid-red: grid cells with ≥ 20 bird and 5 primate species, dark-red: grid cells with ≥ 50 bird and 10 primate species. Key Biodiversity Areas: 1 Itombwe Mountains, 2 Luama Kivu, 3 Kabobo, 4 Lake Nokoué, 5 Afi River, 6 Mount Oku; Protected Areas: A Boé, B Dulombi, C Cantanhez Forest, D Outamba, E Kilimi, F Loma, G Sankan Biriwa, H Comoé, I Nimba, K Digya, L Bomu, M Garamba, N Okapis, O Virunga, P Kahuzi-Biega, Q Kibira, R Cross River.

Both taxa show substantial overlap in the type of habitat used (Figure 5, Table S2). Forest habitats are used by almost all primate species (97%) and represent an important habitat for birds as well (48% of the 218 bird species occurring within primate range), making forests the primary shared habitat between these taxa. But, both groups also occupy artificial terrestrial habitats (65% of birds and 23% of primates), savannas (30% of birds and 12% of primates), and shrubland (57% of birds and 11% of primates). Grasslands, wetlands, and rocky areas are used more extensively by birds than by primates.

**Figure 5.**
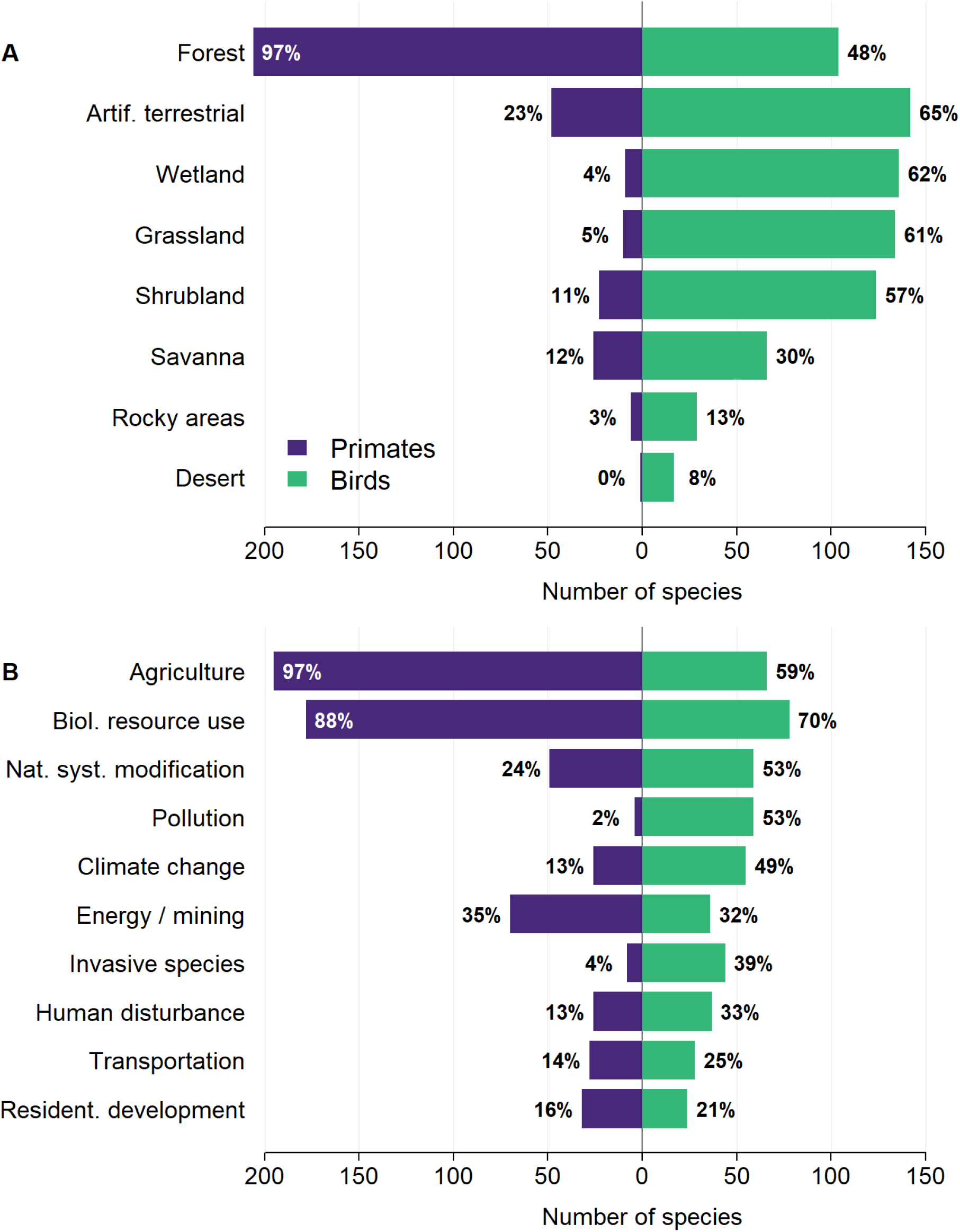
Comparison of (A) habitat use and (B) threats listed by the IUCN Red List of Threatened Species for primates and birds. Bars depict the number of species and percentage values indicate the corresponding percentage of species. Information was available for (A) 212 primate and 218 bird species, and for (B) 202 primate and 112 bird species.

### 3.2 Threats and conservation status

Among the 213 primate species, 156 are classified as Threatened and 18 as Near-threatened (IUCN 2025). 90% of primate species have declining populations. Among the 218 migratory bird species, 43% have declining populations and 24 species are classified as Threatened (Table S3).

For 202 primate species and 112 bird species IUCN lists threat information. Both birds and primates face substantial threats from agriculture (97% primates, 59% birds), and biological resource use (88% primates, 70% birds, Figure 5, Table S4). Mining is the third most significant threat to primate species (35%) and is also listed as a threat to 32% of bird species. The relative importance of other threats differs between the taxa. Pollution and natural system modifications affect a higher proportion of assessed bird species (53% each) compared to primates (2% and 24%, respectively). Also, invasive species and diseases pose a substantially greater threat to birds (39%) than to primates (4%). Climate change and severe weather are listed as threats to 49% of bird species and 13% of primates. Approximately, 54% of protected areas overlap with at least one grid cell of the areas identified of highest diversity in primate and wintering bird species (dark-red areas in Figure 4).

### 3.3 Conservation interventions

The Conservation Evidence Project lists 120 beneficial or likely beneficial interventions for birds, compared to 12 for primates. This is in line with a well-documented evidence gap on the effectiveness of conservation interventions for primates (Junker *et al* 2020).

Of the 120 bird interventions, we rated 101 (84%) interventions as not applicable to primates, as they address biological traits or threats unique to birds (e.g., artificially incubate and hand-rear birds, provision of artificial nesting sites, predator and brood parasite control, Table S5). From the 19 bird interventions applicable to primates, 10 corresponded to 7 unique entries in the primate Conservation Evidence database, of which two have been assessed as ‘likely beneficial’ (implement no-hunting community policies and create/protect habitat corridors), while the other interventions had limited evidence or have not been assessed for primates yet (Junker *et al* 2017). The remaining nine interventions include actions related to habitat restoration, and mitigation of threats such as agricultural intensification and infrastructure development. While these actions are specifically evaluated for birds, their ecological scope suggests they could contribute to primate conservation where species ranges and habitat requirements overlap.

Of the 12 primate interventions, we rated eight (67%) as applicable to birds and they correspond to three unique bird conservation interventions. Two of those have been assessed as (likely) beneficial, specifically, ‘reducing conflict by deterring birds from crops’ and ‘restore or create bird habitat’ (Williams *et al* 2013). One intervention (‘Run research project and ensure permanent human presence at site’) is applicable to birds, but has not been evaluated.

Overall, several interventions show proven benefits for both taxa, specifically those aiming to create, restore and protect habitat, as well as reducing hunting pressure. A number of additional interventions have demonstrated effectiveness for one taxon and are plausibly applicable to the other, but have not yet been evaluated: including interventions related to managing agricultural areas (e.g. pesticide use), mitigating the impact of infrastructure developments, and establishing research sites with permanent human presence.

## 4 Discussion

Our analysis reveals that African primates and Afro-Palearctic migratory birds overlap spatially but also in the habitats they use and threats they face. This co-occurrence, combined with a set of conservation interventions that benefit both taxa, indicates that coordinated conservation management efforts targeting both taxa simultaneously offer potential for conservation synergies. While conserving migratory bird habitat has been found to potentially benefit resident wildlife (Wilson *et al* 2022), our results show that the reverse is also true.

Hunting and agriculture are the two dominant threats affecting both primates and migratory birds. Bushmeat studies across West and Central Africa indicate that primates and medium-sized birds are harvested under similar economic and cultural contexts (Luiselli *et al* 2019, Taylor *et al* 2015). Agricultural expansion, particularly the conversion of land to cocoa, oil palm, and smallholder farms, drives primate habitat loss (Estrada *et al* 2017). Similarly, agricultural expansion degrades bird wintering areas through habitat loss and degradation (Perrings and Halkos 2015, Douglas *et al* 2023).

However, some threats are better documented for one taxon than the other, and cross-taxon research could help close these evidence gaps. For example, the effects of pesticides on birds are relatively well studied, with evidence that they deplete food and nesting resources, reduce breeding success, and cause direct mortality (Moreau *et al* 2022). Their impacts on African primates, by contrast, remain poorly understood. Evidence from Asia shows that agrochemicals in oil-palm systems can reduce infant survival in macaques (Holzner *et al* 2024), suggesting that pesticide exposure may pose comparable risks for African primates in agricultural landscapes.

Similarly, our review of evidence on conservation interventions suggests that habitat protection and reducing hunting pressure could benefit both taxa. For example, protected areas across Africa support both endemic primates and important populations of migratory species (Kaboumba *et al* 2025, Davenport *et al* 2014). However, the evidence-base for conservation intervention effectiveness is small for both taxa. A recent study, for example, showed that none of the studies on interventions for agriculture and birds is located in Africa (González-Suárez *et al* 2026).

Building on these findings, we identify three conservation policy priorities that can operationalize cross-taxon synergies.

### 4.1 Strengthening cross-taxa scientific collaboration and species monitoring

Strengthening scientific collaboration, knowledge exchange and cross-taxa species monitoring would lay the foundation for coordinated conservation action. Permanent primate field stations have been established across large parts of the primate range. Bird wintering habitats, by contrast, remain severely undersurveyed, and the infrastructure needed to monitor them is largely absent (Table 1). Cross-taxa monitoring methods such as passive acoustic monitoring and camera trapping can be deployed cost-effectively within existing primate research structures. At the same time, primate populations in savanna-forest ecotones are relatively less studied, but this is where we found a strong overlap in intermediate primate richness and the highest bird richness. Thus, ornithologists and primatologists would benefit from expanding research infrastructure into these areas.

**Table 1.** Overview of synergies for integrated conservation efforts for African primates and Afro-Palearctic migratory birds.

| Current status for primates | Current status for migratory birds | Conservation synergy | Benefit |
| --- | --- | --- | --- |
| **Strengthening cross-taxa scientific collaboration and species monitoring** |  |  |  |
| Permanent field stations established across primate ranges | No equivalent field infrastructure in African wintering areas | Co-financing of field stations by primate and bird organisations | Primates: improved financial sustainability of existing field stations,<br>Birds: access to established field infrastructure; can serve as hubs for studies on Afro-Palearctic migratory birds |
| Field surveys and monitoring programmes are regularly implemented | Wintering population status under-surveyed | Integration of bird surveys into existing wildlife monitoring schemes; development of joint acoustic monitoring and camera trapping protocols | Both taxa: sharing of complementary knowledge through joint survey planning and data analyses increases quality of outcomes |
| **Improving conservation evidence base for shared threats in African context** |  |  |  |
| Evidence base on interventions well-recognised as severely limited | Existing intervention evidence base drawn mostly from non-African contexts | Co-designed conservation evidence studies at shared field sites applicable to both taxa (e.g., habitat restoration, agricultural mitigation, infrastructure impact reduction, chemical pollution near mining sites) | Both taxa: Africa-specific evidence base generated for interventions |
| Impacts of specific threats not well-documented, e.g. impact of pesticide exposure | Pesticide use well-documented as a threat to birds | Primate researchers adopt pesticide monitoring protocols developed for birds; joint assessment of agrochemical impacts in areas of highest co-occurrence | Primates: new threat category quantified,<br>Birds: expanded spatial coverage of pesticide threat monitoring |
| **Creating synergies in conservation finance** |  |  |  |
| Primate conservation attracts substantial European funding; primates function as flagship and | Wintering habitat in Africa receives negligible dedicated conservation funding | Reframe existing European primate conservation funding to explicitly account for Afro-Palearctic migratory birds through adjusted donor criteria | Primates: increased financial sustainability through diversified donor base |
| umbrella species for broader habitat protection |  | and joint funding bids by primate and bird organisations targeting areas of highest co-occurrence | Birds: dedicated funding for African wintering habitat conservation |
| Great ape presence usually triggers 'Critical Habitat' classification, and companies have to implement mitigation measures (IFC Performance Standard 6) | Migratory birds can trigger 'Critical Habitat' but rarely do so in practice | Include co-occurring migratory birds into ape-triggered mitigation and offset measures | Birds: private sector financed mitigation and offset measures expanded to include bird wintering areas |
| Conservation action plans exist for threatened primate species; implemented through African range-state partnerships | No equivalent conservation action plans addressing wintering habitat for most Afro-Palearctic migrants | Joint conservation action planning explicitly incorporating wintering habitat needs alongside breeding area priorities; remote responsibility framing used to engage European governments and NGOs as accountable stakeholders | Primates: action plans broadened to attract intercontinental co-financing, Birds: wintering habitat needs formally incorporated into species-level conservation planning |

### 4.2 Improving the conservation evidence base for shared threats in the African context

A coordinated approach can improve the conservation evidence base for threats shared by African primates and wintering birds. The monitoring infrastructure described under priority (i) would help to generate systematic, comparable data on species occurrence and threat exposure across sites and over time. In addition, before-after and control-impact approaches can generate the data needed to evaluate what conservation interventions actually work. Shared study designs and integrating research on conservation effectiveness across both taxa simultaneously would reduce overall costs of securing funding, setting up research design, and establishing monitoring systems.

### 4.3 Creating synergies in conservation finance

To translate the above priorities into sustained action can be supported by creating synergies in conservation finance. Funding for conservation across Africa is heavily skewed towards charismatic flagship species, driven by international donor preferences and revenue from photographic tourism and trophy hunting (Lindsey *et al* 2020). For example, Gorilla-based tourism accounted for roughly half of the Uganda Wildlife Authority’s annual budget (Tumusiime and Vedeld 2012). Thus, financing mechanisms focused on primates could be modified to explicitly incorporate incentives for protecting bird wintering habitats within the same landscapes.

In private development finance (e.g., for mining sites), the presence of great apes usually results in an area being classified as ‘Critical habitat’ (CH) and then mitigation measures reducing ecological damage have to be implemented by the company (International Finance Corporation’s Performance Standard 6, Junker *et al* 2024). Though migratory species can also trigger CH, they rarely do in practice (Serckx *et al* 2018). Incorporating migratory birds into ape-focused mitigation measures would extend these investments to co-occurring birds at low additional costs.

Conversely, existing frameworks for migratory species conservation already contain provisions that could extend protection to co-occurring taxa. The Convention on Migratory Species, for example, includes mechanisms to encourage protection of non-listed species that share habitat with listed ones. This is illustrated by the Migratory Soaring Birds Project, funded by the Global Environment Facility and implemented by BirdLife International (GEF 2022). The project integrated bird conservation into five industry sectors along the Rift Valley/Red Sea flyway, including banning toxic pesticides in agriculture, modifying wind turbines and power lines to reduce electrocution risk, and improving waste management. With this the project achieved conservation outcomes that extended beyond migratory birds. Thus, designing financing instruments that formally recognise the co-occurrence of African primates and migratory birds would allow conservation investments to generate returns across both taxa simultaneously.

In conclusion, these three priorities outline a path toward coordinated conservation based on existing infrastructure and funding mechanisms. Given that Afro-Palearctic migratory birds receive disproportionately little conservation attention across their African wintering areas, integrated policies across both taxa offer conservation synergies that neither group achieves when managed in isolation.

## Acknowledgments

Substantial part of this work emerged from the workshop “Training great ape range country nationals in wildlife survey data analysis, interpretation and paper writing” that took place in Rwanda (2024). We are grateful to the following organizations which organized the training: the Senckenberg Museum of Natural History Görlitz (Germany), the Dian Fossey Gorilla Fund in Kinigi—DFGF (Rwanda), the IUCN SSC Primate Specialist Group—Section on Great Apes (PSG-SGA), Re:wild (USA) and the African Primatological Society (APS). We particularly thank Prof. Inza Kone and Dr. Winnie Eckardt for their special investment in the success of this workshop which enabled the collaborative work for this research.

## Data availability statement

Species range data are available from the IUCN Red List of Threatened Species (IUCN 2025) and BirdLife International (http://datazone.birdlife.org). Species occurrence data are available from GBIF database (Calabuig 2013). Information on species threat status, threats and habitat are available from the IUCN Red List of Threatened Species. R-code for the analysis will be made available via Zenodo upon publication.

