## Supporting Information for "Birds protected in Europe, overlooked in Africa: conservation synergies between African primates and wintering Afro-Palearctic migratory birds"

### Supplementary tables

Table S1 | Proportion of overlap of geographic range

Table S2 | Habitat type

Table S3 | Conservation status and population trends

Table S4 | Threat

Table S5 | Conservation interventions beneficial for birds

Table S6 | Conservation interventions beneficial for primates

### Supplementary figures

Figure S1 | Distribution of the minimum distance between bird–primate occurrences

**Table S1** | Overlap of geographic range for each combination of bird and primate species with at least 30% overlap.

| **Bird species** | **Bird range [1000 km2]** | **Primate species** | **Primate range [1000 km2]** | **Overlap [1000 km2]** | **Overlap [%]** |
|---|---|---|---|---|---|
| Luscinia luscinia | 1696.2 | Galago moholi | 4126.9 | 1661.0 | 97.92 |
| Emberiza hortulana | 1005.8 | Galago senegalensis | 7752.8 | 891.9 | 88.68 |
| Lanius minor | 2086.2 | Papio ursinus | 3347.9 | 1759.9 | 84.36 |
| Streptopelia turtur | 4529.9 | Galago senegalensis | 7752.8 | 3772.9 | 83.29 |
| Botaurus stellaris | 2973.2 | Papio anubis | 7813.3 | 2460.0 | 82.74 |
| Botaurus stellaris | 2973.2 | Galago senegalensis | 7752.8 | 2449.4 | 82.38 |
| Jynx torquilla | 6042.2 | Papio anubis | 7813.3 | 4964.4 | 82.16 |
| Luscinia luscinia | 1696.2 | Otolemur crassicaudatus | 4843.0 | 1386.4 | 81.74 |
| Podiceps cristatus | 1156.8 | Papio ursinus | 3347.9 | 939.2 | 81.19 |
| Botaurus stellaris | 2973.2 | Erythrocebus patas | 6601.8 | 2351.9 | 79.1 |
| Circaetus gallicus | 5408.5 | Galago senegalensis | 7752.8 | 4140.8 | 76.56 |
| Merops apiaster | 2204.9 | Galago moholi | 4126.9 | 1680.9 | 76.23 |
| Jynx torquilla | 6042.2 | Galago senegalensis | 7752.8 | 4599 | 76.11 |
| Luscinia megarhynchos | 4728.7 | Papio anubis | 7813.3 | 3552.7 | 75.13 |
| Luscinia megarhynchos | 4728.7 | Galago senegalensis | 7752.8 | 3463.4 | 73.24 |
| Tachymarptis melba | 5852.6 | Papio anubis | 7813.3 | 4252 | 72.65 |
| Saxicola rubetra | 8316.6 | Galago senegalensis | 7752.8 | 5996.2 | 72.1 |
| Streptopelia turtur | 4529.9 | Papio anubis | 7813.3 | 3256.4 | 71.89 |
| Lymnocryptes minimus | 8897.7 | Papio anubis | 7813.3 | 6362.1 | 71.5 |
| Lymnocryptes minimus | 8897.7 | Galago senegalensis | 7752.8 | 6360 | 71.48 |
| Botaurus stellaris | 2973.2 | Chlorocebus tantalus | 3941.9 | 2124.7 | 71.46 |
| Streptopelia turtur | 4529.9 | Erythrocebus patas | 6601.8 | 3220.7 | 71.1 |
| Accipiter nisus | 2707.2 | Papio anubis | 7813.3 | 1906.4 | 70.42 |
| Platalea leucorodia | 547.4 | Galago senegalensis | 7752.8 | 384.8 | 70.3 |
| Lymnocryptes minimus | 8897.7 | Erythrocebus patas | 6601.8 | 6216.5 | 69.87 |
| Saxicola rubetra | 8316.6 | Papio anubis | 7813.3 | 5801.9 | 69.76 |
| Merops apiaster | 2204.9 | Otolemur crassicaudatus | 4843 | 1537.4 | 69.73 |
| Anthus cervinus | 7783.5 | Papio anubis | 7813.3 | 5425.7 | 69.71 |
| Circaetus gallicus | 5408.5 | Papio anubis | 7813.3 | 3722.4 | 68.82 |
| Luscinia megarhynchos | 4728.7 | Erythrocebus patas | 6601.8 | 3244.3 | 68.61 |
| Jynx torquilla | 6042.2 | Erythrocebus patas | 6601.8 | 4145 | 68.6 |
| Glareola pratincola | 8002.5 | Galago senegalensis | 7752.8 | 5487.6 | 68.57 |
| Tringa erythropus | 9895.3 | Papio anubis | 7813.3 | 6763 | 68.35 |
| Calidris temminckii | 9982.2 | Galago senegalensis | 7752.8 | 6753.6 | 67.66 |
| Anthus cervinus | 7783.5 | Galago senegalensis | 7752.8 | 5259.1 | 67.57 |
| Apus pallidus | 7010 | Galago senegalensis | 7752.8 | 4728.5 | 67.45 |
| Tachymarptis melba | 5852.6 | Galago senegalensis | 7752.8 | 3936.4 | 67.26 |
| Spatula querquedula | 9682.6 | Galago senegalensis | 7752.8 | 6493.6 | 67.06 |
| Anas acuta | 8775 | Galago senegalensis | 7752.8 | 5823.7 | 66.37 |

| | | | | | |
|---|---|---|---|---|---|
| Charadrius alexandrinus | 4947.6 | Galago senegalensis | 7752.8 | 3269.9 | 66.09 |
| Accipiter nisus | 2707.2 | Galago senegalensis | 7752.8 | 1786.6 | 65.99 |
| Oenanthe oenanthe | 10064.2 | Galago senegalensis | 7752.8 | 6628.5 | 65.86 |
| Calidris temminckii | 9982.2 | Papio anubis | 7813.3 | 6530.7 | 65.42 |
| Tringa erythropus | 9895.3 | Galago senegalensis | 7752.8 | 6459.4 | 65.28 |
| Cecropis daurica | 508.1 | Papio anubis | 7813.3 | 330.1 | 64.97 |
| Spatula querquedula | 9682.6 | Papio anubis | 7813.3 | 6189.6 | 63.92 |
| Oenanthe pleschanka | 3244 | Papio anubis | 7813.3 | 2066.6 | 63.71 |
| Oenanthe pleschanka | 3244 | Galago senegalensis | 7752.8 | 2040 | 62.89 |
| Emberiza hortulana | 1005.8 | Papio anubis | 7813.3 | 632.1 | 62.85 |
| Charadrius dubius | 10419 | Papio anubis | 7813.3 | 6523 | 62.61 |
| Anas acuta | 8775 | Papio anubis | 7813.3 | 5475.7 | 62.4 |
| Cecropis daurica | 508.1 | Erythrocebus patas | 6601.8 | 316.1 | 62.21 |
| Platalea leucorodia | 547.4 | Erythrocebus patas | 6601.8 | 340 | 62.11 |
| Lanius excubitor | 1212.8 | Galago senegalensis | 7752.8 | 743.3 | 61.29 |
| Apus pallidus | 7010 | Papio anubis | 7813.3 | 4289.6 | 61.19 |
| Circaetus gallicus | 5408.5 | Erythrocebus patas | 6601.8 | 3299.1 | 61 |
| Apus pallidus | 7010 | Erythrocebus patas | 6601.8 | 4249.6 | 60.62 |
| Recurvirostra avosetta | 4242.1 | Galago senegalensis | 7752.8 | 2568.7 | 60.55 |
| Spatula clypeata | 4872.8 | Galago senegalensis | 7752.8 | 2948.3 | 60.51 |
| Glareola pratincola | 8002.5 | Papio anubis | 7813.3 | 4780 | 59.73 |
| Riparia riparia | 11235.3 | Papio anubis | 7813.3 | 6645.4 | 59.15 |
| Charadrius dubius | 10419 | Galago senegalensis | 7752.8 | 6101.9 | 58.57 |
| Accipiter brevipes | 2408.2 | Papio anubis | 7813.3 | 1385.8 | 57.55 |
| Merops apiaster | 2204.9 | Chlorocebus cynosuros | 3084.9 | 1255.8 | 56.95 |
| Oenanthe oenanthe | 10064.2 | Papio anubis | 7813.3 | 5720.5 | 56.84 |
| Gallinago gallinago | 13637.7 | Galago senegalensis | 7752.8 | 7750.7 | 56.83 |
| Calidris temminckii | 9982.2 | Erythrocebus patas | 6601.8 | 5661.8 | 56.72 |
| Egretta gularis | 102.3 | Galago senegalensis | 7752.8 | 58 | 56.7 |
| Cecropis daurica | 508.1 | Galago senegalensis | 7752.8 | 288 | 56.68 |
| Riparia riparia | 11235.3 | Galago senegalensis | 7752.8 | 6338.7 | 56.42 |
| Gallinago gallinago | 13637.7 | Papio anubis | 7813.3 | 7682.1 | 56.33 |
| Ciconia nigra | 7011.1 | Papio anubis | 7813.3 | 3946.9 | 56.3 |
| Charadrius alexandrinus | 4947.6 | Papio anubis | 7813.3 | 2783.4 | 56.26 |
| Larus fuscus | 7978.8 | Papio anubis | 7813.3 | 4474.3 | 56.08 |
| Ciconia nigra | 7011.1 | Galago senegalensis | 7752.8 | 3905.3 | 55.7 |
| Egretta gularis | 102.3 | Chlorocebus aethiops | 1145.3 | 56.7 | 55.43 |
| Aythya fuligula | 4073.3 | Papio anubis | 7813.3 | 2254.3 | 55.34 |
| Monticola saxatilis | 5818.2 | Galago senegalensis | 7752.8 | 3205 | 55.09 |
| Saxicola rubetra | 8316.6 | Erythrocebus patas | 6601.8 | 4533.8 | 54.52 |
| Luscinia luscinia | 1696.2 | Chlorocebus pygerythrus | 4650.1 | 924.6 | 54.51 |
| Aythya fuligula | 4073.3 | Galago senegalensis | 7752.8 | 2218.2 | 54.46 |
| Recurvirostra avosetta | 4242.1 | Erythrocebus patas | 6601.8 | 2309 | 54.43 |

| | | | | | |
|---|---|---|---|---|---|
| Gelochelidon nilotica | 6164.5 | Galago senegalensis | 7752.8 | 3354.2 | 54.41 |
| Spatula clypeata | 4872.8 | Papio anubis | 7813.3 | 2637.5 | 54.13 |
| Glareola pratincola | 8002.5 | Erythrocebus patas | 6601.8 | 4312.2 | 53.89 |
| Circus aeruginosus | 12992.7 | Galago senegalensis | 7752.8 | 6999.5 | 53.87 |
| Anas crecca | 6213.6 | Galago senegalensis | 7752.8 | 3342.7 | 53.8 |
| Ficedula semitorquata | 4987.9 | Papio anubis | 7813.3 | 2634.4 | 52.82 |
| Coturnix coturnix | 10558.6 | Galago senegalensis | 7752.8 | 5542.9 | 52.5 |
| Tachymarptis melba | 5852.6 | Erythrocebus patas | 6601.8 | 3068.5 | 52.43 |
| Mareca penelope | 5155.3 | Papio anubis | 7813.3 | 2693.7 | 52.25 |
| Tringa erythropus | 9895.3 | Erythrocebus patas | 6601.8 | 5151.5 | 52.06 |
| Spatula querquedula | 9682.6 | Erythrocebus patas | 6601.8 | 5037.3 | 52.02 |
| Sylvia nisoria | 2329.5 | Papio anubis | 7813.3 | 1196.3 | 51.35 |
| Anthus cervinus | 7783.5 | Erythrocebus patas | 6601.8 | 3996.1 | 51.34 |
| Buteo buteo | 8570.1 | Otolemur crassicaudatus | 4843 | 4375.2 | 51.05 |
| Monticola saxatilis | 5818.2 | Papio anubis | 7813.3 | 2962.2 | 50.91 |
| Tringa totanus | 10755.4 | Galago senegalensis | 7752.8 | 5440.8 | 50.59 |
| Podiceps cristatus | 1156.8 | Chlorocebus pygerythrus | 4650.1 | 583.2 | 50.41 |
| Tringa totanus | 10755.4 | Papio anubis | 7813.3 | 5416.8 | 50.36 |
| Charadrius alexandrinus | 4947.6 | Erythrocebus patas | 6601.8 | 2482.3 | 50.17 |
| Luscinia megarhynchos | 4728.7 | Chlorocebus tantalus | 3941.9 | 2360.4 | 49.92 |
| Oenanthe oenanthe | 10064.2 | Erythrocebus patas | 6601.8 | 5016.5 | 49.84 |
| Jynx torquilla | 6042.2 | Chlorocebus tantalus | 3941.9 | 3007.6 | 49.78 |
| Anas crecca | 6213.6 | Papio anubis | 7813.3 | 3090.7 | 49.74 |
| Clanga pomarina | 7643.8 | Otolemur crassicaudatus | 4843 | 3778.2 | 49.43 |
| Cecropis daurica | 508.1 | Chlorocebus aethiops | 1145.3 | 251 | 49.4 |
| Acrocephalus palustris | 6535.7 | Chlorocebus pygerythrus | 4650.1 | 3225.3 | 49.35 |
| Gyps fulvus | 5263.3 | Galago senegalensis | 7752.8 | 2596.9 | 49.34 |
| Platalea leucorodia | 547.4 | Papio anubis | 7813.3 | 269.5 | 49.23 |
| Ficedula semitorquata | 4987.9 | Galago senegalensis | 7752.8 | 2453.6 | 49.19 |
| Circus aeruginosus | 12992.7 | Papio anubis | 7813.3 | 6379.7 | 49.1 |
| Accipiter nisus | 2707.2 | Erythrocebus patas | 6601.8 | 1329 | 49.09 |
| Sylvia curruca | 5242.8 | Galago senegalensis | 7752.8 | 2570.9 | 49.04 |
| Larus fuscus | 7978.8 | Galago senegalensis | 7752.8 | 3902.2 | 48.91 |
| Sylvia nisoria | 2329.5 | Galago senegalensis | 7752.8 | 1129.3 | 48.48 |
| Lanius excubitor | 1212.8 | Papio anubis | 7813.3 | 583 | 48.07 |
| Coturnix coturnix | 10558.6 | Papio anubis | 7813.3 | 5075.8 | 48.07 |
| Anas acuta | 8775 | Erythrocebus patas | 6601.8 | 4205.4 | 47.92 |
| Emberiza hortulana | 1005.8 | Erythrocebus patas | 6601.8 | 481.3 | 47.85 |
| Mareca penelope | 5155.3 | Galago senegalensis | 7752.8 | 2464.6 | 47.81 |
| Recurvirostra avosetta | 4242.1 | Papio anubis | 7813.3 | 2017.8 | 47.57 |
| Coturnix coturnix | 10558.6 | Erythrocebus patas | 6601.8 | 5016.1 | 47.51 |
| Egretta gularis | 102.3 | Papio anubis | 7813.3 | 48 | 46.92 |
| Oenanthe isabellina | 6904 | Galago senegalensis | 7752.8 | 3232.6 | 46.82 |

| | | | | | |
|---|---|---|---|---|---|
| Tringa totanus | 10755.4 | Erythrocebus patas | 6601.8 | 5003 | 46.52 |
| Gallinago gallinago | 13637.7 | Erythrocebus patas | 6601.8 | 6318.6 | 46.33 |
| Circus aeruginosus | 12992.7 | Erythrocebus patas | 6601.8 | 6008.3 | 46.24 |
| Gelochelidon nilotica | 6164.5 | Papio anubis | 7813.3 | 2837.9 | 46.04 |
| Ficedula albicollis | 6278.9 | Papio anubis | 7813.3 | 2883 | 45.92 |
| Luscinia luscinia | 1696.2 | Chlorocebus cynosuros | 3084.9 | 776.1 | 45.76 |
| Buteo buteo | 8570.1 | Galago moholi | 4126.9 | 3914.9 | 45.68 |
| Circus pygargus | 14167.1 | Galago senegalensis | 7752.8 | 6451.6 | 45.54 |
| Pelecanus onocrotalus | 14258.1 | Galago senegalensis | 7752.8 | 6435.3 | 45.13 |
| Anthropoides virgo | 1626.6 | Papio anubis | 7813.3 | 733.1 | 45.07 |
| Charadrius asiaticus | 9251.4 | Chlorocebus pygerythrus | 4650.1 | 4169.8 | 45.07 |
| Caprimulgus europaeus | 10503 | Galago senegalensis | 7752.8 | 4712.7 | 44.87 |
| Accipiter brevipes | 2408.2 | Galago senegalensis | 7752.8 | 1075.6 | 44.66 |
| Buteo buteo | 8570.1 | Chlorocebus pygerythrus | 4650.1 | 3811.5 | 44.47 |
| Charadrius dubius | 10419 | Erythrocebus patas | 6601.8 | 4616.1 | 44.3 |
| Oenanthe isabellina | 6904 | Papio anubis | 7813.3 | 3050.1 | 44.18 |
| Hieraaetus pennatus | 13919 | Galago senegalensis | 7752.8 | 6057 | 43.52 |
| Iduna pallida | 9253 | Papio anubis | 7813.3 | 4022.5 | 43.47 |
| Clanga clanga | 531.7 | Papio anubis | 7813.3 | 230.3 | 43.31 |
| Riparia riparia | 11235.3 | Erythrocebus patas | 6601.8 | 4864.9 | 43.3 |
| Ficedula albicollis | 6278.9 | Galago senegalensis | 7752.8 | 2716.8 | 43.27 |
| Crex crex | 9624.5 | Chlorocebus pygerythrus | 4650.1 | 4158.6 | 43.21 |
| Iduna pallida | 9253 | Galago senegalensis | 7752.8 | 3989.6 | 43.12 |
| Sylvia nisoria | 2329.5 | Chlorocebus aethiops | 1145.3 | 998.5 | 42.86 |
| Luscinia luscinia | 1696.2 | Papio ursinus | 3347.9 | 722.1 | 42.57 |
| Aquila nipalensis | 9291.4 | Otolemur crassicaudatus | 4843 | 3935.8 | 42.36 |
| Clanga pomarina | 7643.8 | Chlorocebus pygerythrus | 4650.1 | 3232.1 | 42.28 |
| Lymnocryptes minimus | 8897.7 | Chlorocebus tantalus | 3941.9 | 3745.3 | 42.09 |
| Merops apiaster | 2204.9 | Papio kindae | 2213 | 927.4 | 42.06 |
| Clanga pomarina | 7643.8 | Galago moholi | 4126.9 | 3213.4 | 42.04 |
| Calidris pugnax | 17940.1 | Galago senegalensis | 7752.8 | 7497.9 | 41.79 |
| Lanius nubicus | 5293.2 | Galago senegalensis | 7752.8 | 2206.3 | 41.68 |
| Motacilla cinerea | 2140.6 | Galago senegalensis | 7752.8 | 886.6 | 41.42 |
| Sylvia curruca | 5242.8 | Erythrocebus patas | 6601.8 | 2161.5 | 41.23 |
| Tringa ochropus | 18810.7 | Galago senegalensis | 7752.8 | 7745.2 | 41.17 |
| Lanius minor | 2086.2 | Chlorocebus pygerythrus | 4650.1 | 858.1 | 41.13 |
| Motacilla alba | 9066.1 | Galago senegalensis | 7752.8 | 3723.6 | 41.07 |
| Tringa ochropus | 18810.7 | Papio anubis | 7813.3 | 7700.9 | 40.94 |
| Luscinia luscinia | 1696.2 | Cercopithecus mitis | 2347.9 | 693.3 | 40.87 |
| Emberiza hortulana | 1005.8 | Chlorocebus aethiops | 1145.3 | 410.6 | 40.82 |
| Pelecanus onocrotalus | 14258.1 | Papio anubis | 7813.3 | 5748.1 | 40.31 |
| Caprimulgus europaeus | 10503 | Papio anubis | 7813.3 | 4228.5 | 40.26 |
| Hippolais polyglotta | 8866.8 | Erythrocebus patas | 6601.8 | 3555.9 | 40.1 |

| | | | | | |
|---|---|---|---|---|---|
| Aythya ferina | 4189.1 | Papio anubis | 7813.3 | 1672 | 39.91 |
| Aythya ferina | 4189.1 | Galago senegalensis | 7752.8 | 1668.5 | 39.83 |
| Larus fuscus | 7978.8 | Erythrocebus patas | 6601.8 | 3175.1 | 39.79 |
| Neophron percnopterus | 10823.4 | Galago senegalensis | 7752.8 | 4302.6 | 39.75 |
| Apus apus | 18335.5 | Papio anubis | 7813.3 | 7285.9 | 39.74 |
| Otus scops | 17405.2 | Papio anubis | 7813.3 | 6906.6 | 39.68 |
| Gyps fulvus | 5263.3 | Erythrocebus patas | 6601.8 | 2081.8 | 39.55 |
| Hieraaetus pennatus | 13919 | Papio anubis | 7813.3 | 5504.3 | 39.55 |
| Sylvia curruca | 5242.8 | Papio anubis | 7813.3 | 2063.2 | 39.35 |
| Neophron percnopterus | 10823.4 | Erythrocebus patas | 6601.8 | 4248.6 | 39.25 |
| Calidris pugnax | 17940.1 | Papio anubis | 7813.3 | 7039.7 | 39.24 |
| Tringa erythropus | 9895.3 | Chlorocebus tantalus | 3941.9 | 3873.4 | 39.14 |
| Cuculus canorus | 12118.4 | Otolemur crassicaudatus | 4843 | 4727.7 | 39.01 |
| Lanius minor | 2086.2 | Galago moholi | 4126.9 | 813.9 | 39.01 |
| Anthropoides virgo | 1626.6 | Galago senegalensis | 7752.8 | 634.1 | 38.98 |
| Apus apus | 18335.5 | Galago senegalensis | 7752.8 | 7129.8 | 38.89 |
| Circus pygargus | 14167.1 | Papio anubis | 7813.3 | 5508.8 | 38.88 |
| Motacilla alba | 9066.1 | Papio anubis | 7813.3 | 3515.6 | 38.78 |
| Larus ridibundus | 1180.5 | Galago senegalensis | 7752.8 | 457.6 | 38.76 |
| Spatula clypeata | 4872.8 | Erythrocebus patas | 6601.8 | 1887 | 38.73 |
| Otus scops | 17405.2 | Galago senegalensis | 7752.8 | 6732.6 | 38.68 |
| Oenanthe pleschanka | 3244 | Chlorocebus pygerythrus | 4650.1 | 1252.3 | 38.6 |
| Aquila nipalensis | 9291.4 | Chlorocebus pygerythrus | 4650.1 | 3574.2 | 38.47 |
| Lanius senator | 16511.5 | Erythrocebus patas | 6601.8 | 6300.1 | 38.16 |
| Buteo buteo | 8570.1 | Papio ursinus | 3347.9 | 3270.4 | 38.16 |
| Xenus cinereus | 1022.1 | Chlorocebus pygerythrus | 4650.1 | 387.8 | 37.94 |
| Clanga clanga | 531.7 | Galago senegalensis | 7752.8 | 201.2 | 37.84 |
| Otus scops | 17405.2 | Erythrocebus patas | 6601.8 | 6575 | 37.78 |
| Aquila nipalensis | 9291.4 | Galago moholi | 4126.9 | 3509.5 | 37.77 |
| Buteo rufinus | 4128.2 | Galago senegalensis | 7752.8 | 1555.7 | 37.68 |
| Ciconia ciconia | 17342.4 | Galago senegalensis | 7752.8 | 6512.2 | 37.55 |
| Luscinia luscinia | 1696.2 | Papio kindae | 2213 | 635.9 | 37.49 |
| Larus ridibundus | 1180.5 | Papio anubis | 7813.3 | 442 | 37.44 |
| Lanius nubicus | 5293.2 | Erythrocebus patas | 6601.8 | 1979.1 | 37.39 |
| Gyps fulvus | 5263.3 | Papio anubis | 7813.3 | 1967.2 | 37.38 |
| Crex crex | 9624.5 | Otolemur crassicaudatus | 4843 | 3585.8 | 37.26 |
| Iduna opaca | 7164.1 | Erythrocebus patas | 6601.8 | 2663.3 | 37.18 |
| Chlidonias hybrida | 17455 | Erythrocebus patas | 6601.8 | 6487.3 | 37.17 |
| Ficedula hypoleuca | 14573 | Erythrocebus patas | 6601.8 | 5414.5 | 37.15 |
| Charadrius asiaticus | 9251.4 | Otolemur crassicaudatus | 4843 | 3432.9 | 37.11 |
| Ptyonoprogne rupestris | 975.3 | Chlorocebus aethiops | 1145.3 | 360.7 | 36.98 |
| Tringa glareola | 20967.8 | Galago senegalensis | 7752.8 | 7746 | 36.94 |
| Chlidonias hybrida | 17455 | Papio anubis | 7813.3 | 6440 | 36.89 |

| | | | | | |
|---|---|---|---|---|---|
| Saxicola rubetra | 8316.6 | Chlorocebus tantalus | 3941.9 | 3067.5 | 36.88 |
| Tringa glareola | 20967.8 | Papio anubis | 7813.3 | 7700.3 | 36.72 |
| Caprimulgus europaeus | 10503 | Chlorocebus pygerythrus | 4650.1 | 3842 | 36.58 |
| Muscicapa striata | 20967.1 | Papio anubis | 7813.3 | 7608.8 | 36.29 |
| Muscicapa striata | 20967.1 | Galago senegalensis | 7752.8 | 7596.8 | 36.23 |
| Phylloscopus sibilatrix | 17616.8 | Erythrocebus patas | 6601.8 | 6373.9 | 36.18 |
| Apus pallidus | 7010 | Chlorocebus tantalus | 3941.9 | 2529.9 | 36.09 |
| Buteo rufinus | 4128.2 | Papio anubis | 7813.3 | 1489.5 | 36.08 |
| Lanius senator | 16511.5 | Galago senegalensis | 7752.8 | 5946.5 | 36.01 |
| Cuculus canorus | 12118.4 | Galagoides thomasi | 4397 | 4359.3 | 35.97 |
| Hydroprogne caspia | 3260.2 | Papio anubis | 7813.3 | 1172.3 | 35.96 |
| Oenanthe cypriaca | 1560.4 | Papio anubis | 7813.3 | 558.9 | 35.82 |
| Cuculus canorus | 12118.4 | Galagoides demidoff | 4339.1 | 4337 | 35.79 |
| Tringa nebularia | 21712.8 | Galago senegalensis | 7752.8 | 7739.4 | 35.64 |
| Tringa nebularia | 21712.8 | Papio anubis | 7813.3 | 7700.3 | 35.46 |
| Lanius nubicus | 5293.2 | Papio anubis | 7813.3 | 1876.7 | 35.45 |
| Phoenicurus phoenicurus | 17150.5 | Galago senegalensis | 7752.8 | 6058 | 35.32 |
| Pandion haliaetus | 21964.6 | Galago senegalensis | 7752.8 | 7737.3 | 35.23 |
| Tringa stagnatilis | 19573.5 | Papio anubis | 7813.3 | 6890 | 35.2 |
| Neophron percnopterus | 10823.4 | Papio anubis | 7813.3 | 3803.6 | 35.14 |
| Ptyonoprogne rupestris | 975.3 | Papio anubis | 7813.3 | 342 | 35.07 |
| Pandion haliaetus | 21964.6 | Papio anubis | 7813.3 | 7700.3 | 35.06 |
| Actitis hypoleucos | 22123.9 | Galago senegalensis | 7752.8 | 7749.7 | 35.03 |
| Ficedula albicollis | 6278.9 | Chlorocebus pygerythrus | 4650.1 | 2198.2 | 35.01 |
| Glareola nordmanni | 5222.1 | Papio anubis | 7813.3 | 1827.4 | 34.99 |
| Limosa limosa | 20191 | Galago senegalensis | 7752.8 | 7062.3 | 34.98 |
| Lanius senator | 16511.5 | Papio anubis | 7813.3 | 5768.2 | 34.93 |
| Accipiter nisus | 2707.2 | Chlorocebus tantalus | 3941.9 | 945.7 | 34.93 |
| Ptyonoprogne rupestris | 975.3 | Galago senegalensis | 7752.8 | 339.8 | 34.84 |
| Monticola solitarius | 1965.4 | Galago senegalensis | 7752.8 | 684.6 | 34.83 |
| Actitis hypoleucos | 22123.9 | Papio anubis | 7813.3 | 7700.3 | 34.81 |
| Lanius excubitor | 1212.8 | Erythrocebus patas | 6601.8 | 421.9 | 34.79 |
| Oenanthe pleschanka | 3244 | Chlorocebus aethiops | 1145.3 | 1127.9 | 34.77 |
| Gallinago media | 21077.5 | Papio anubis | 7813.3 | 7316.9 | 34.71 |
| Motacilla cinerea | 2140.6 | Papio anubis | 7813.3 | 742 | 34.66 |
| Chlidonias leucopterus | 22365.5 | Galago senegalensis | 7752.8 | 7722.6 | 34.53 |
| Chlidonias leucopterus | 22365.5 | Papio anubis | 7813.3 | 7718.6 | 34.51 |
| Ciconia ciconia | 17342.4 | Papio anubis | 7813.3 | 5980.5 | 34.48 |
| Porzana porzana | 11276.2 | Chlorocebus pygerythrus | 4650.1 | 3888.5 | 34.48 |
| Aquila heliaca | 1683.2 | Papio anubis | 7813.3 | 579.9 | 34.45 |
| Hippolais olivetorum | 12180.1 | Chlorocebus pygerythrus | 4650.1 | 4192.9 | 34.42 |
| Tringa ochropus | 18810.7 | Erythrocebus patas | 6601.8 | 6475.2 | 34.42 |
| Mareca strepera | 3518.4 | Galago senegalensis | 7752.8 | 1209.6 | 34.38 |

| | | | | | |
|---|---|---|---|---|---|
| Circus macrourus | 19600.5 | Galago senegalensis | 7752.8 | 6736.7 | 34.37 |
| Oenanthe isabellina | 6904 | Erythrocebus patas | 6601.8 | 2370.9 | 34.34 |
| Ciconia nigra | 7011.1 | Erythrocebus patas | 6601.8 | 2403.6 | 34.28 |
| Gallinago media | 21077.5 | Galago senegalensis | 7752.8 | 7222.6 | 34.27 |
| Chlidonias hybrida | 17455 | Galago senegalensis | 7752.8 | 5972.4 | 34.22 |
| Hippolais polyglotta | 8866.8 | Galago senegalensis | 7752.8 | 3032.6 | 34.2 |
| Hirundo rustica | 14159.4 | Otolemur crassicaudatus | 4843 | 4842.6 | 34.2 |
| Phylloscopus sibilatrix | 17616.8 | Papio anubis | 7813.3 | 6015.2 | 34.14 |
| Hieraaetus pennatus | 13919 | Erythrocebus patas | 6601.8 | 4749.5 | 34.12 |
| Luscinia megarhynchos | 4728.7 | Colobus guereza | 2595.3 | 1609.5 | 34.04 |
| Anthus cervinus | 7783.5 | Chlorocebus tantalus | 3941.9 | 2644.9 | 33.98 |
| Anas crecca | 6213.6 | Erythrocebus patas | 6601.8 | 2109.3 | 33.95 |
| Acrocephalus scirpaceus | 19160.3 | Papio anubis | 7813.3 | 6499.7 | 33.92 |
| Tringa stagnatilis | 19573.5 | Galago senegalensis | 7752.8 | 6633.2 | 33.89 |
| Limosa limosa | 20191 | Papio anubis | 7813.3 | 6824.9 | 33.8 |
| Accipiter brevipes | 2408.2 | Erythrocebus patas | 6601.8 | 809.1 | 33.6 |
| Phoenicurus phoenicurus | 17150.5 | Papio anubis | 7813.3 | 5757.1 | 33.57 |
| Coracias garrulus | 18969.1 | Galago senegalensis | 7752.8 | 6363.9 | 33.55 |
| Pelecanus onocrotalus | 14258.1 | Erythrocebus patas | 6601.8 | 4764.9 | 33.42 |
| Motacilla cinerea | 2140.6 | Chlorocebus aethiops | 1145.3 | 711.7 | 33.25 |
| Luscinia megarhynchos | 4728.7 | Galagoides thomasi | 4397 | 1571.2 | 33.23 |
| Sylvia atricapilla | 12162.4 | Galago senegalensis | 7752.8 | 4038.9 | 33.21 |
| Circaetus gallicus | 5408.5 | Chlorocebus tantalus | 3941.9 | 1794 | 33.17 |
| Mareca strepera | 3518.4 | Papio anubis | 7813.3 | 1165.3 | 33.12 |
| Calidris pugnax | 17940.1 | Erythrocebus patas | 6601.8 | 5928.4 | 33.05 |
| Larus fuscus | 7978.8 | Chlorocebus tantalus | 3941.9 | 2635.1 | 33.03 |
| Cuculus canorus | 12118.4 | Papio anubis | 7813.3 | 3998.1 | 32.99 |
| Oenanthe cypriaca | 1560.4 | Galago senegalensis | 7752.8 | 514 | 32.94 |
| Acrocephalus scirpaceus | 19160.3 | Galago senegalensis | 7752.8 | 6308.1 | 32.92 |
| Hirundo rustica | 14159.4 | Chlorocebus pygerythrus | 4650.1 | 4648.7 | 32.83 |
| Calidris temminckii | 9982.2 | Chlorocebus tantalus | 3941.9 | 3273.5 | 32.79 |
| Anthus trivialis | 23634.1 | Galago senegalensis | 7752.8 | 7722.4 | 32.67 |
| Tadorna ferruginea | 596.9 | Chlorocebus aethiops | 1145.3 | 195 | 32.67 |
| Falco vespertinus | 20760.2 | Papio anubis | 7813.3 | 6779.4 | 32.66 |
| Ficedula albicollis | 6278.9 | Otolemur crassicaudatus | 4843 | 2044.3 | 32.56 |
| Phylloscopus sibilatrix | 17616.8 | Galago senegalensis | 7752.8 | 5717 | 32.45 |
| Anthus trivialis | 23634.1 | Papio anubis | 7813.3 | 7649.2 | 32.37 |
| Clanga pomarina | 7643.8 | Galago senegalensis | 7752.8 | 2467.5 | 32.28 |
| Acrocephalus palustris | 6535.7 | Otolemur crassicaudatus | 4843 | 2109.1 | 32.27 |
| Phoenicurus phoenicurus | 17150.5 | Erythrocebus patas | 6601.8 | 5523.8 | 32.21 |
| Aquila nipalensis | 9291.4 | Galago senegalensis | 7752.8 | 2991.6 | 32.2 |
| Acrocephalus scirpaceus | 19160.3 | Erythrocebus patas | 6601.8 | 6165.3 | 32.18 |
| Pernis apivorus | 18941.1 | Papio anubis | 7813.3 | 6093.6 | 32.17 |

| | | | | | |
|---|---|---|---|---|---|
| Aquila heliaca | 1683.2 | Galago senegalensis | 7752.8 | 541 | 32.14 |
| Phylloscopus bonelli | 9082 | Erythrocebus patas | 6601.8 | 2910.4 | 32.05 |
| Cuculus canorus | 12118.4 | Galago moholi | 4126.9 | 3879.3 | 32.01 |
| Circus pygargus | 14167.1 | Erythrocebus patas | 6601.8 | 4535.3 | 32.01 |
| Merops apiaster | 2204.9 | Cercopithecus mitis | 2347.9 | 703.9 | 31.92 |
| Tachymarptis melba | 5852.6 | Chlorocebus pygerythrus | 4650.1 | 1858.2 | 31.75 |
| Riparia riparia | 11235.3 | Chlorocebus tantalus | 3941.9 | 3564 | 31.72 |
| Jynx torquilla | 6042.2 | Colobus guereza | 2595.3 | 1914.3 | 31.68 |
| Locustella fluviatilis | 13264.4 | Papio anubis | 7813.3 | 4201.8 | 31.68 |
| Charadrius asiaticus | 9251.4 | Galago moholi | 4126.9 | 2922.6 | 31.59 |
| Clanga pomarina | 7643.8 | Papio anubis | 7813.3 | 2406.8 | 31.49 |
| Hippolais olivetorum | 12180.1 | Papio anubis | 7813.3 | 3823.2 | 31.39 |
| Calidris falcinellus | 94.7 | Papio hamadryas | 458.3 | 29.7 | 31.36 |
| Hippolais icterina | 23411.5 | Papio anubis | 7813.3 | 7339.5 | 31.35 |
| Cuculus canorus | 12118.4 | Chlorocebus pygerythrus | 4650.1 | 3771.1 | 31.12 |
| Gallinago media | 21077.5 | Erythrocebus patas | 6601.8 | 6554.6 | 31.1 |
| Charadrius dubius | 10419 | Chlorocebus tantalus | 3941.9 | 3219.4 | 30.9 |
| Iduna opaca | 7164.1 | Galago senegalensis | 7752.8 | 2212.4 | 30.88 |
| Coracias garrulus | 18969.1 | Papio anubis | 7813.3 | 5853.7 | 30.86 |
| Falco naumanni | 21067 | Galago senegalensis | 7752.8 | 6502.1 | 30.86 |
| Egretta gularis | 102.3 | Papio hamadryas | 458.3 | 31.5 | 30.79 |
| Tachymarptis melba | 5852.6 | Chlorocebus tantalus | 3941.9 | 1797.6 | 30.71 |
| Sylvia communis | 19631.4 | Galago senegalensis | 7752.8 | 6013.9 | 30.63 |
| Pernis apivorus | 18941.1 | Galago senegalensis | 7752.8 | 5784.1 | 30.54 |
| Glareola pratincola | 8002.5 | Chlorocebus tantalus | 3941.9 | 2433.4 | 30.41 |
| Merops apiaster | 2204.9 | Papio ursinus | 3347.9 | 670.2 | 30.4 |
| Charadrius dubius | 10419 | Galagoides thomasi | 4397 | 3167.2 | 30.4 |
| Circus macrourus | 19600.5 | Papio anubis | 7813.3 | 5958.6 | 30.4 |
| Aquila heliaca | 1683.2 | Chlorocebus aethiops | 1145.3 | 510.9 | 30.35 |
| Spatula querquedula | 9682.6 | Chlorocebus tantalus | 3941.9 | 2938.8 | 30.35 |
| Locustella luscinioides | 1937 | Galago senegalensis | 7752.8 | 587.4 | 30.33 |
| Falco vespertinus | 20760.2 | Galago senegalensis | 7752.8 | 6297.3 | 30.33 |
| Numenius phaeopus | 3711.4 | Papio anubis | 7813.3 | 1125.3 | 30.32 |
| Crex crex | 9624.5 | Galago senegalensis | 7752.8 | 2912.5 | 30.26 |
| Hydroprogne caspia | 3260.2 | Galago senegalensis | 7752.8 | 983.8 | 30.18 |
| Gelochelidon nilotica | 6164.5 | Chlorocebus pygerythrus | 4650.1 | 1850 | 30.01 |

**Table S2.** Habitat type for each primate species (n=212, habitat info not available for one species) and each bird species occurring within primate range (n=218) compiled from the IUCN Red List of Threatened Species (IUCN 2025). Typically, several types of habitats are listed for each species.

| Habitat | No. of primate species | % of primate species | No. of bird species | % of bird species |
|---|---|---|---|---|
| Artificial aquatic marine | 0 | 0 | 62 | 28 |
| Artificial terrestrial | 48 | 23 | 142 | 65 |
| Caves | 2 | 1 | 0 | 0 |
| Desert | 1 | 0 | 17 | 8 |
| Forest | 206 | 97 | 104 | 48 |
| Grassland | 10 | 5 | 134 | 61 |
| Marine coastal supratidal | 0 | 0 | 52 | 24 |
| Marine intertidal | 0 | 0 | 63 | 29 |
| Marine neritic | 0 | 0 | 42 | 19 |
| Marine oceanic | 0 | 0 | 2 | 1 |
| Rocky areas | 6 | 3 | 29 | 13 |
| Savanna | 26 | 12 | 66 | 30 |
| Shrubland | 23 | 11 | 124 | 57 |
| Wetland | 9 | 4 | 136 | 62 |

**Table S3.** Conservation status and population trends for each primate species (n=213) and bird species occurring within primate range (n=218) compiled from the IUCN Red List of Threatened Species (IUCN 2025). CR = Critically Endangered; EN = Endangered; VU = Vulnerable, NT = Near threatened, LC = Least concern, DD = Data deficient.

| Taxa | Conservation status | | | | Population trend | | | |
|---|---|---|---|---|---|---|---|---|
| | Threatened (CR/EN/VU) | NT | LC | DD | Declining | Stable | Increasing | Unknown |
| Primates | 156 | 18 | 36 | 3 | 190 | 11 | 1 | 11 |
| Birds | 9 | 15 | 194 | 0 | 94 | 64 | 36 | 24 |

**Table S4.** Threat type for each primate species (n=202) and bird species (n=112) for which threats were assessed by the IUCN Red List of Threatened Species (IUCN 2025). Typically, several types of threats are listed for each species.

| Threat | No. of primate species | % of primate species | No. of bird species | % of bird species |
|---|---|---|---|---|
| Agriculture | 195 | 97 | 66 | 59 |
| Biological resource use | 178 | 88 | 78 | 70 |

| | | | | |
|---|---|---|---|---|
| Climate change / severe weather | 26 | 13 | 55 | 49 |
| Energy production / mining | 70 | 35 | 36 | 32 |
| Human intrusions / disturbance | 26 | 13 | 37 | 33 |
| Invasive species, genes, diseases | 8 | 4 | 44 | 39 |
| Natural system modifications | 49 | 24 | 59 | 53 |
| Pollution | 4 | 2 | 59 | 53 |
| Residential / commercial development | 32 | 16 | 24 | 21 |
| Transportation service corridors | 28 | 14 | 28 | 25 |

**Table S5.** List of 120 conservation interventions listed as (likely) beneficial by Conservation Evidence (CE) (Williams *et al* 2013), including attribution to high-level category according to CE, whether intervention is applicable to primates, conservation interventions that are similar for primates and their effectiveness rating according to CE. See separate file "Table_S5_bird_conservation_interventions.xlsx".

**Table S6.** List of 12 conservation interventions listed as (likely) beneficial by Conservation Evidence (Junker *et al* 2017), including attribution to high-level category according to CE, whether intervention is applicable to birds, conservation interventions that are similar for birds and their effectiveness rating according to CE. See separate file "Table_S6_primate_conservation_interventions.xlsx".

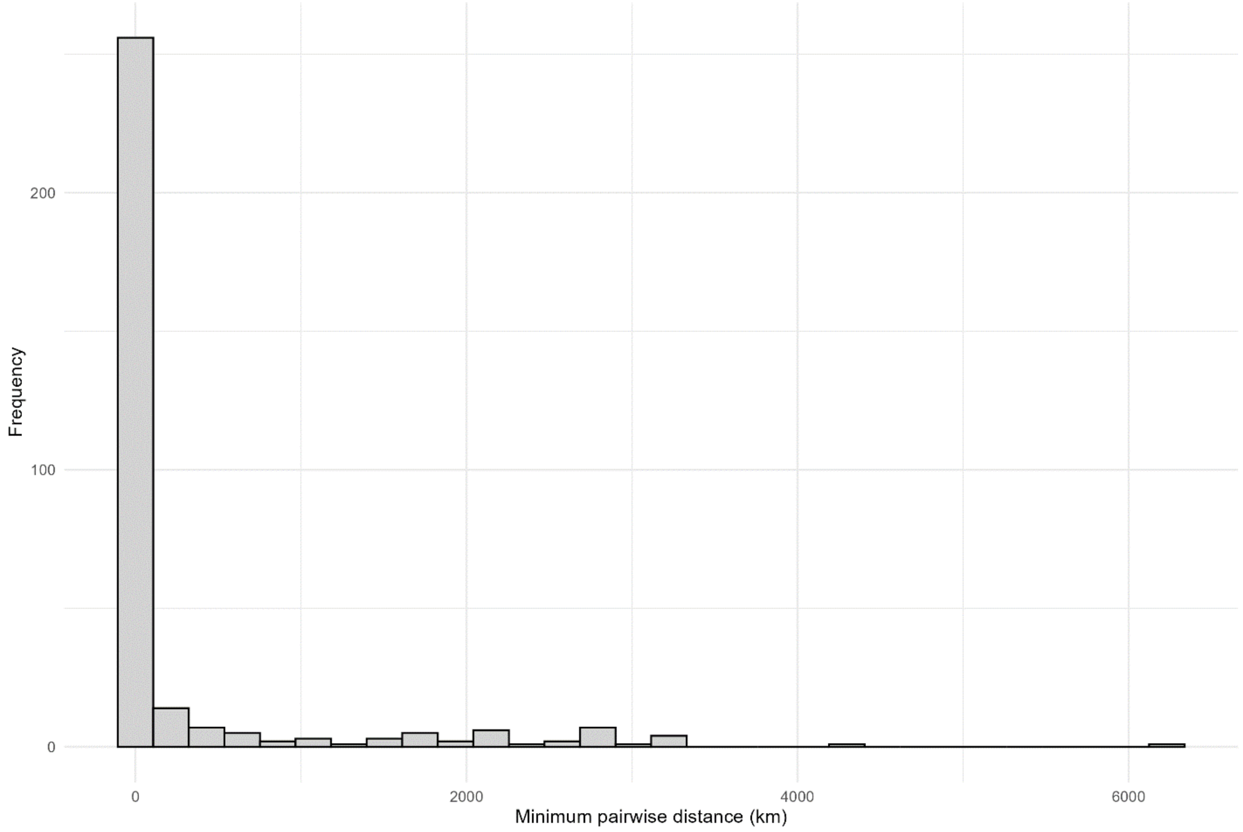


**Figure S1.** Minimum distance of occurrence points of primate – bird species pairs for which spatial ranges overlapped by more than 80% and for which occurrence data was available from GBIF (n = 332 primate – bird species pairs).
